# High-resolution perfusion imaging in rodents using pCASL at 9.4 T

**DOI:** 10.1101/2024.03.07.583839

**Authors:** Sara Pires Monteiro, Lydiane Hirschler, Emmanuel L. Barbier, Patricia Figueiredo, Noam Shemesh

## Abstract

Adequate perfusion is critical for maintaining normal brain function and aberrations thereof are hallmarks of many diseases. Pseudo-Continuous Arterial Spin Labeling (pCASL) MRI enables noninvasive perfusion mapping without contrast agent injection and with higher signal-to-noise ratio (SNR) than alternative methods. Despite its great potential, pCASL remains challenging, unstable, and relatively low-resolution in rodents – especially in mice – thereby limiting the investigation of perfusion properties in many transgenic or other relevant rodent models of disease. Here, we address this gap by developing a novel experimental setup for high-resolution pCASL imaging in mice and combining it with the enhanced SNR of cryogenic probes. We show that our new experimental setup allows for optimal positioning of the carotids within the cryogenic coil, rendering labelling reproducible. With the proposed methodology, we managed to increase the spatial resolution of pCASL perfusion images by a factor of 15 in mice; a factor of 6 in rats is gained compared to the state of the art just by virtue of the cryogenic coil. We also show that the improved pCASL perfusion imaging allows much better delineation of specific brain areas, both in healthy animals as well as in rat and mouse models of stroke. Our results bode well for future high-definition pCASL perfusion imaging in rodents.

**Data availability:** Data will be freely made available upon request.

## 1. Introduction

Cerebral Blood Flow (CBF), or perfusion, is critical for supplying nutrients and oxygen to the brain, thereby maintaining normal metabolism and function.^1,2^ CBF quantification is typically achieved using Arterial Spin Labeling (ASL) – a non-invasive Magnetic Resonance Imaging (MRI) technique harnessing spin-labeled arterial water as an endogenous tracer.^3,4^ ASL is often used in clinical settings due to its advantageous avoidance of contrast agent injection^5^, which also allows repeated measurements, and its delivery of quantitative CBF measurements in absolute units of ml of blood per 100 g of tissue per minute.^6^

ASL can be categorized into three main labeling strategies: Continuous ASL (CASL), pseudo continuous ASL (pCASL) and Pulsed ASL (PASL). In PASL, labeling is performed by applying a brief radiofrequency (RF) inversion pulse over a wide region (labeling slab) in the neck region.^7^ In CASL, a higher signal-to-noise ratio (SNR) is achieved by instead employing an RF pulse to a thinner labeling plane, typically using a dedicated RF coil over the neck region, for a relatively extended period of time to produce flow-driven adiabatic inversion. On the other hand, pCASL replaces the continuous pulse by a train of shorter RF pulses, thereby avoiding the need for a second RF coil^8^, albeit at the cost of a somewhat lower labeling efficiency.^6^ In practice, due to its relative advantages, pCASL is often preferred over both CASL and PASL.

In pre-clinical settings, small rodents such as rats and mice provide ample opportunities for research due to the ease of interrogating biological processes under controlled conditions, and further due to the existence of many transgenic or other (e.g. lesion) models of disease. To date however, pCASL MRI remains very challenging in such preclinical settings, and is limited to relatively low resolution imaging.^9–13^ The application of pCASL in rodents is complicated by multiple factors, when compared to humans, including geometric constraints such as smaller brain size and higher blood velocity. Combined with the typical use of higher field strengths, these factors lead to strong off-resonance effects that severely affect the pCASL inversion efficiency (α).^13,14^ B1 field inhomogeneities at the labeling plane are also potentially exacerbating factors since more power is needed to reach the same flip angle in an inhomogeneous field. This leads to higher power deposition and consequently higher specific absorption rate (SAR).^8^

Hirschler *et al*.^15^ recently implemented a pCASL sequence for perfusion imaging in rats at 9.4 T using a room temperature coil. To deal with the high B_0_ inhomogeneities at the labeling plane, a prescan was introduced to allow for the optimization of the phase corrections of the pCASL pulse train. This optimization step proved crucial to achieve sufficiently high α in the rat. With the optimized corrections, CBF maps could be obtained with state-of-the art resolution of 234x234x1000 μm^3^. Later, the pCASL sequence was also implemented at 7T for perfusion imaging in mice, again with a room temperature coil, yielding multi-slice CBF images at a resolution of 225x225x1500 μm^3^.^9,10,16^ The authors also implemented a 3D-pCASL-EPI sequence in combination with a cryogenic coil in mice and used stronger gradients to deal with the reduced inversion efficiency.^17^ More recently, the pCASL sequence along with the respective phase correction scheme was successfully adjusted to assess blood brain barrier permeability in mice at 11.7 T.^18^ Still, in these studies, large variability in and CBF were reported, and the spatial resolution remained relatively low for many rodent neuroimaging applications. High resolution and more robust pCASL would clearly benefit a detailed representation of small brain structures as well as one’s ability to assess brain alterations and brain injury at earlier stages of disease^19,20^. We note in passing that other approaches for mapping surrogates of CBF with high resolution exist, but they are typically indirect and localized in nature.^21–23^

Here, we set out to develop a methodology for significantly increasing the resolution of pCASL perfusion images at 9.4 Tesla, in rats as well as in mice. We harness a cryogenic coil^24^ that provides high SNR, together with Hirschler et al.’s optimized pCASL sequence^15^ that is robust against B_0_ inhomogeneities.^16^ We first demonstrated high-resolution pCASL perfusion imaging in rats, and then introduce a novel experimental setup for mouse perfusion imaging. The setup is required for optimal positioning of the carotids for pCASL labeling in the mice, and we show that it dramatically improves α and reproducibility of pCASL imaging. These improvements allowed us to improve the spatial resolution by over an order of magnitude compared to the state of the art in mice. We further demonstrate the utility of high resolution pCASL imaging in mouse and rat models of stroke, where we also achieved a significant improvement in spatial resolution.

## 2. Methods

All animal experiments were preapproved by the competent institutional and national authorities and carried out according to European Directive 2010/63.

### 2.1. Animals

For the mouse experiments, N=16 adult female C57BL/g mice were used (∼ 11.1±0.6 weeks old, weights 22.5±1.5, grown with a 12 h light/dark cycle with ad libitum access to food and water). In the rat experiments, N=4 Long-Evans female rats, 7.3±1.3 weeks old, weight: 245.6 56.7g were used.

### 2.2. Mouse and rat stroke model: surgical Procedures

In 3 mice (out of the N=16) and 1 rat (out of the N=4), a photothrombotic stroke model^25^ was used to induce a cerebral infarction in the somatosensory cortex (S1). Mice were administered with buprenorphine Mouse Mix (1:20 dilution of Bupaq (0,3 mg/ml) in NaCl/Water, 100 *μ*l/mouse, 15-30 g body weight, subcutaneous) 30 min prior to surgery. Their body temperature was kept constant at 37°C with the use of an electrical heating pad. The mice were anesthetized with 1.5-2.5% isoflurane. The stroke was induced by injecting a solution of Rose Bengal dye (Sigma Aldrich, Portugal) (10 mg/ml) which was delivered intravenously by retroorbital injection (10 l/g body weight). The animals were subsequently irradiated with a cold light source (beam light intensity of 5 W/cm^2^) in the left S1 (1.07 mm posterior and 1.5 mm lateral to bregma^26^) for 15 minutes. The 3 mice were longitudinally assessed at 3 different timepoints: immediately after stroke (t=0h), one day after stroke (t=24h) and 5 days after stroke (t= 5d). In one rat, the procedure was repeated in right S1 (1.08 mm posterior and 3.5 mm lateral to bregma^27^): dye was delivered intravenously (13 mg/kg body weight) through tail vein injection. The rat was scanned 1 day after stroke.

### 2.3. MRI Experiments

#### 2.3.1. Experimental Setup

Experiments were conducted on a 9.4 T Bruker Biospec Scanner (Karlsruhe, Germany) operated by an AVANCE III HD console, equipped with an 86 mm volume coil for transmittance, a 4-element array mouse or rat cryogenic coil for signal reception and a gradient system able to generate up to 660 mT/m isotropically. The temperature of the cryogenic coil heater was set at 37°C.

A custom-built stereolithography 3D printed resin *ramp* (Figure 1, panel B) was designed for the mouse scans to prevent the bending of the animal’s head relative to the body caused by presence of the ear bars and mouth bite bar – *ramp setup*. Design files for the *ramp* can be found on github (https://github.com/sarapmonteiro/ramp) and in Supplementary Information. For the rats, no *ramp* was required due to the larger anatomy of the animals that does not require the use of ear bars to prevent head-motion.

**Figure 1.**
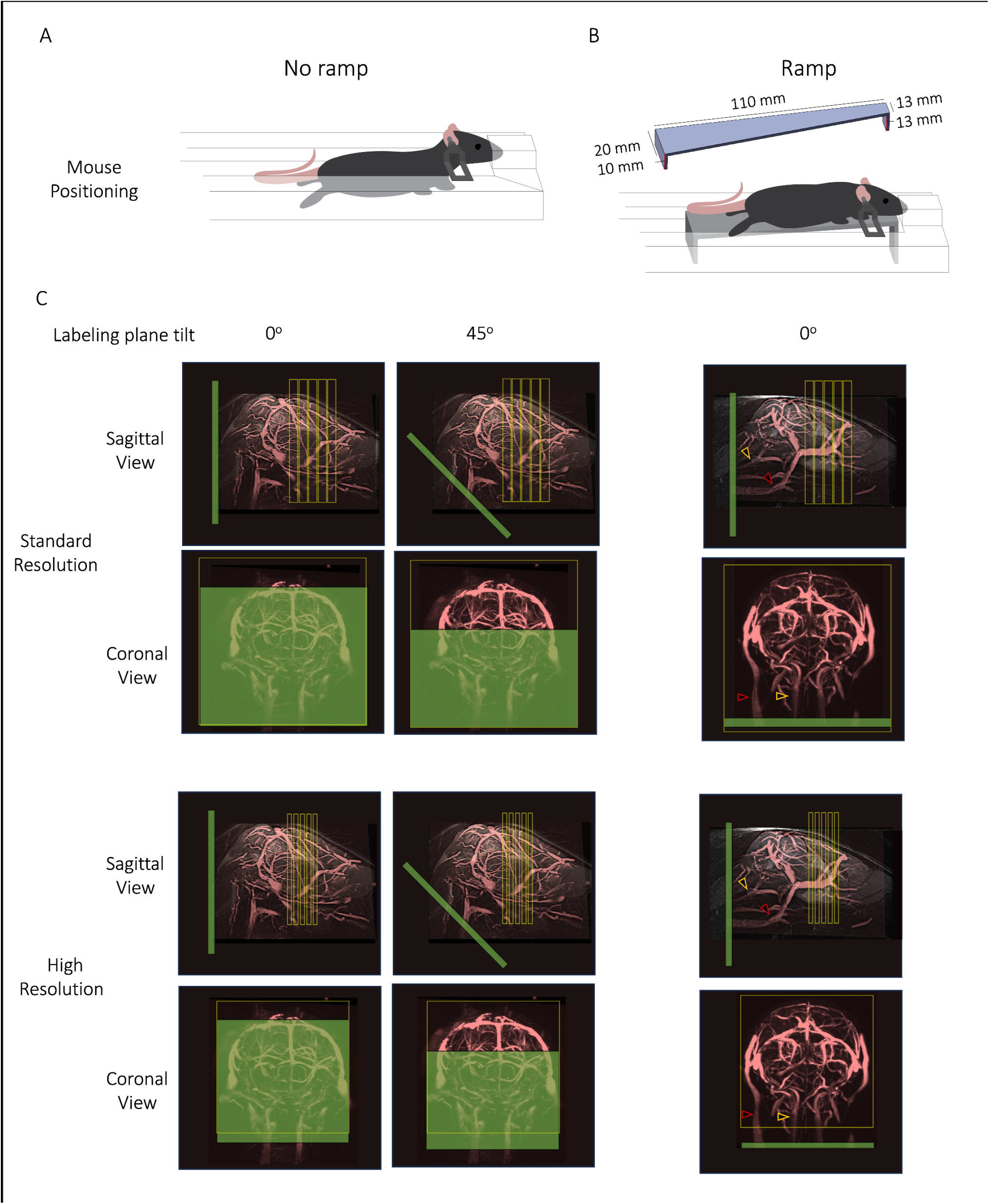
Illustration of the labelling plane and slice prescription in both setups and resolutions. Setups for mouse positioning in the scanner when using a cryogenic coil for no ramp (A) and ramp (B); C) Placement of labeling and imaging regions concerning the brain and major vessels irrigating the brain, overlayed on an anatomical sagittal midline image and a maximum intensity projection of time-of-flight angiography. Left) *No ramp setup*: labeling plane tilt of 0° (*no ramp 0°*) and 45° (*no ramp 45°*) in relation to the imaging region; In the *no ramp 0μ* the labeling plane is parallel to both the main feeding arteries and the imaging region, appearing to be superimposed in the coronal view. In the *no ramp 45°* the labeling plane angle appears to be insufficient to be perpendicular to the carotids, due to their extreme bending. Still, it overlaps the imaging region in the coronal perspective. Right) *Ramp setup*: labeling plane angle set to 0° and mice positioned on top of a *ramp*. Animals were scanned at standard and high-resolution. The labeling plane is perpendicular to both the main feeding arteries and parallel to the imaging region. Note that a relatively large FOV localizer was necessary to assess the accurate location of the carotids. Carotid and vertebral arteries are highlighted by the yellow and red arrowheads, respectively.

#### 2.3.2. Animal Preparation

Anaesthesia was induced via a mixture of medical air and 3.5% isoflurane (Vetflurane, Virbac, France) until they were still, with no response to strong foot squeeze or righting reflex. During this process, the body temperature of the animals was kept constant using an electrical heating pad. Prior to moving the mouse to the scanner, the isoflurane concentration was reduced to 2.5%, and then kept between 2.5%-1.5% for maintenance. Oxygen concentration was regulated with an oxygen monitor (MX300-I, Viamed, United Kingdom) and maintained at 50%. The mouse was weighed and transferred to the scanner bed.

Eye ointment (Bepanthen Eye Drops, Bepanthen, Germany) was applied to avoid damaging the corneas. Temperature was monitored with the use of a rectal probe and maintained at 37°C using a closed-loop temperature regulated water circulation system. Respiratory rate was evaluated using a respiratory sensor (Model 1030 Monitoring Gating System, SAII, United States of America) and kept between 40 and 90 breaths per minute throughout all experiments by adjusting the isoflurane dose. Cardiac metrics were not measured.

Two different setups were tested: the conventional setup (*no ramp*) and the new setup (*ramp*), as described here. A cohort of N=3 rats were positioned prone in the rat bed on top of a heated water pad, their heads were fixed with a bite bar and no ear bars were used (*no ramp)*. N=7 mice were positioned in the same way on top of a mouse bed, with the addition of ear bars to properly fix their heads (*no ramp*) (Figure 1, panel A). Another group of animals (N=6, all mice) were prepared as mentioned above, but they were further placed above the custom-built *ramp* (Figure 1, panel B).

#### 2.3.3. pCASL Acquisition

An unbalanced pCASL sequence^28^ was used as described in Hirschler *et al*. (2018)^15^. The inversion was achieved through a train of Hanning window-shaped pulses: 400 μs duration, 800 μs interpulse delay, B_1_ of 5μT, G_max_/G_ave_ of 45/5 mT/m, where G_max_ is the gradient applied during the RF pulse.

The interpulse phase offsets were adjusted before the pCASL perfusion measurement using a 2 pre-scan (label and control) scheme to extract the optimal labeling phases^15^. The theoretical phase increment between pulses during the control condition is shifted by 180 degrees compared to that in the label condition to prevent labeling. For the pCASL acquisitions, the labeling duration (LD) was set to 3 s followed by a 300 ms post-labeling delay (PLD). The other parameters were set specifically for mice and rats, as described below.

For the acquisitions in mice, animals not positioned on top of the *ramp* were scanned with the labeling plane tilt of 0° (*no ramp 0*°) and a tilt of 45° (*no ramp 45*o) in relation to the imaging region. The animals previously positioned on top of a *ramp* were scanned with a labeling plane angle of 0° (*ramp*) since their positioning was already perpendicular to the carotid and vertebral arteries (Figure 1, C Right). The labeling plane was positioned at the mouse neck (∼8 mm caudal to the isocenter). For the phase optimization pre-scans, the LD was 1.5 s followed by a 200 ms PLD, and the adjustments were calculated in a single slice positioned in the center of the brain (slice thickness = 2 mm, FOV= 14×14 mm^2^, matrix =96 96, resulting in a spatial resolution of 146×146 *μ*m^2^). The pCASL acquisitions in mice were performed one slice at a time. A total of 5 slices were acquired, separated by a gap of 0.2 mm (standard resolution) or 0.35 mm (high resolution). The following parameters were used:

- Standard resolution pCASL: single-shot spin-echo EPI, FOV= 14×14 mm^2^, slice thickness=1 mm, matrix= 96×96 resulting in an in plane spatial resolution of 146×146 *μ*m^2^, voxel volume=0.021 mm^3^, repetition time (TR)/echo time (TE)= 4000/17 ms, 30 control-label repetitions, T_acq_= 4 min.
- High resolution pCASL: single-shot spin-echo EPI, FOV=12×12 mm^2^, slice thickness= 0.5 mm, matrix= 120×120 resulting in an in plane spatial resolution of 100×100 m^2^, voxel volume= 0.005 mm^3^, TR/TE= 4000/25 ms, 30 control-label repetitions, T_acq_ = 4 min.

For the acquisitions in rats, the *no ramp* setup was used, and the labeling plane was positioned at the rat neck (∼1.3cm caudal to the isocenter). The interpulse phase offsets were set as previously described.^15^ Rat pCASL acquisitions were multi-slice. The pCASL acquisitions were performed with the following parameters:

- Standard resolution pCASL: single-shot spin-echo EPI, FOV=22×22mm^2^, slice thickness=1 mm, matrix=94×94, in plane spatial resolution of 234×234 *μ*m^2^, voxel volume=0.055 mm^3^, TR/TE=4000/40 ms, 4 averages, 30 control-label repetitions, Tacq= 16 min.
- High resolution pCASL: single-shot spin-echo EPI, FOV=19.2×22.0 mm^2^, slice thickness=0.75 mm, matrix= 174 × 200, in plane spatial resolution of 110 × 110 m^2^, voxel volume=0.009 mm^3^, TR/TE=4000/40 ms, 4 averages, 30 control-label repetitions, Tacq= 16 min.

For CBF quantification, a T1 map of the tissue is required. It was obtained from an inversion recovery spin-echo EPI sequence (TR/ TE = 10 000/19 ms; 18 inversion times (TI) between 30 and 10 000 ms; T_acq_ = 4 min) using an adiabatic full passage shaped pulse, generated using a Shinar Leroux algorithm (bandwidth=5000 Hz, length=13.56 ms).^15^ For both mice and rats, a separate T1 map was obtained for each of the pCASL parameter settings, to match the different slice thickness and spatial resolution of both the standard and high-resolution acquisitions. For mice specifically, the T1 maps were also acquired separately for the no *ramp* and the *ramp setup*.

In addition, a pCASL flow-compensated fast low angle shot (fc-FLASH) sequence was employed to estimate the α, by acquiring the signal 3 mm rostral to the labeling plane (1 repetition, TR/TE = 225/5.6 ms; slice thickness= 1 mm, in plane spatial resolution of 172×172 *μ*m^2^, matrix size of 128×128, 2 averages, PLD= 0 ms, LD= 200 ms, T_acq_ = 1m 44 s).15

#### 2.3.4. Angiogram acquisition

In one mouse, to assess the orientation of the labelling plane with respect to the brain vasculature, a 2D time-of-flight (TOF) FLASH angiography sequence was used along with a brain midline sagittal anatomical image with the same FOV. The following parameters were used for the TOF angiography: TR/TE = 15/1.7 ms, 240 slices, slice thickness= 2mm, in plane spatial resolution of 80 × 80 m^2^, 2 averages, FOV=16 16 mm^2^, T_acq_ = 24 min. Anatomicals: T2 RARE, TR/TE = 4000/45 ms, slice thickness = 0.425mm, in plane spatial resolution of 80×80 μm^2^, FOV=16×16 mm^2^, Tacq = 2 min 24 s.

### 2.4. Data analysis

#### 2.4.1. SNR calculation

For each animal, experimental setup, and resolution, the SNR of the raw control pCASL images was calculated voxelwise and in two regions of interest (ROIs) (thalamus and cortex) manually drawn in the raw data of the control series according to the atlas reference^26^ by dividing the mean of the signal across the 30 control repetitions over the standard deviation of the signal across the 30 control repetitions in each voxel (which includes contributions not only from sample and instrument noise but also eddy current effects, physiological instability, etc).

#### 2.4.2. Inversion efficiency

For each animal and experimental setup, the inversion efficiency (*α*) was obtained, for each carotid (left and right), from a complex reconstruction of the pCASL-prepared fc-FLASH, as:

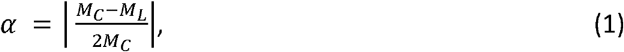

where *M*_*C*_ and *M*_*L*_ are the complex signals from the control and label scans, respectively, averaged across the manually drawn ROIs of the left and right carotids. The final value of α was computed as the mean of the *α* values from the two carotids.

#### 2.4.3. T1 map

For each animal, experimental setup (*ramp/no ramp*), and resolution, a T1 map was obtained by fitting the longitudinal relaxation equation to the voxelwise signal obtained from an inversion recovery spin-echo EPI sequence using a Levenberg-Marquardt algorithm:

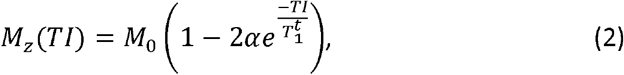

where *M*_*Z*_ is the longitudinal magnetization at each inversion time (*TI*), *M*_0_ is the equilibrium magnetization, 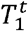 is the longitudinal relaxation time of tissue, and *α* is set to the value from Eq. 1.

#### 2.4.4. rASL map

For each animal, experimental setup, and resolution, the relative ASL (rASL) signal was calculated voxelwise as follows:

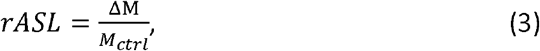

where ΔM is the difference in magnitude between control and label images, averaged across 30 repetitions, and *M_ctrl_* is the magnitude signal across control acquisition averaged across 30 repetitions.

#### 2.4.5. CBF maps

For each animal, experimental setup, and resolution, CBF maps (ml/100g/min)^6,29^ were calculated voxel-by-voxel:

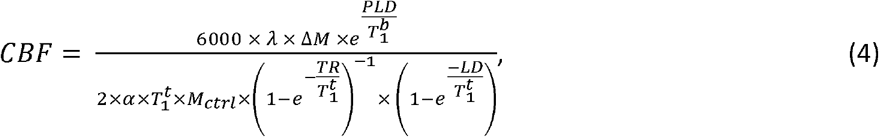

where 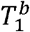 and 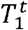 are the longitudinal relaxation times of blood (assumed to be 2430ms at 9.4 T^30^) and tissue (extracted from the T1 map in each voxel), respectively, LD and PLD are the values used in each experiment and *α* is a fixed value obtained from the inversion efficiency fit in Eq. 1. The equilibrium magnetization of blood can be approximated by 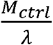, where λ is the blood-tissue partition coefficient of water (the brain average value is considered, 0.9mL/g^31^). The analysis pipeline is presented in Supplementary Figure 1.

#### 2.4.6. Group Average maps

For each experimental setup and resolution, the CBF maps were aligned between animals in ITK-Snap^32^ using only linear and rigid transformations and the median and inter quartile range (IQR) were calculated voxelwise.

#### 2.4.7. ROI Analysis

In the rASL and CBF high-resolution maps of each animal, three ROIs were defined: 1) cortical - right and 2) thalamic – right and 3) left. Averages of the estimated parameters were obtained in each animal, for each setup and for each ROI. To assess inter-hemispheric differences in the rASL and CBF high-resolution maps, the asymmetry index (AI) was calculated in thalamic ROI (left VS right) as follows:

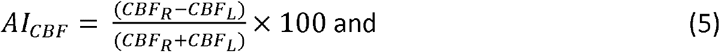

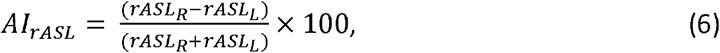

where CBF_R_ and CBF_L_ stand for the CBF values on the right and left sides, respectively and rASL_R_ and rASL_L_ stand for the rASL values on the right and left sides, respectively.

## 3. Results

### 3.1. Mouse pCASL: Raw data and SNR

Harnessing a powerful mouse cryogenic coil, we implemented a mouse setup for pCASL, which required specific adaptations including the *ramp* experimental setup (Figure 1, B). Supplementary Figure 2 presents raw data from a representative mouse for each setup used in this study (*no ramp 0°*, *no ramp 45° and ramp*), both for standard and high-resolution acquisitions. In all cases, the raw data are consistent and with high quality, revealing that all the EPI acquisitions were properly achieved and did not vary within setups. In Supplementary Figure 3, SNR maps of the control images are presented for one representative mouse and analysed across different setups and different resolutions in two ROIs. The SNR distributions across all animals are also presented in Supplementary Figure 3, showing that, as expected, there is greater SNR in all the standard resolution acquisitions when compared to the high resolution maps. In the *ramp* setup, there is systematically greater SNR when compared to other setups. Supplementary Figure 4 shows T1 maps for standard and high-resolution acquisitions for 3 representative animals scanned with the *ramp* and for 3 representative animals scanned with no *ramp* (*0°* and *45°*). The maps present high reproducibility across acquisition settings.

### 3.2. Mouse pCASL: Inversion Efficiency

The results of the inversion efficiency,*α*, calculated for all mice and for each setup (*no ramp 0°*, *no ramp 45° and ramp*) as the average of both carotids, are presented in Figure 2, panel A. The no *ramp 0°* resulted in a of *α* 57.1 ± 20.0%, the *no ramp 45°* resulted in a *α* of 80.2 ± 63.4% and the *ramp* resulted in a *α* of 78.2 ± 4.2%. Note that beyond the improvement in average inversion efficiency, dramatically smaller variance across animals was measured when using the *ramp* setup compared to all other conditions, representing much better reproducibility when using the *ramp* setup.

**Figure 2.**
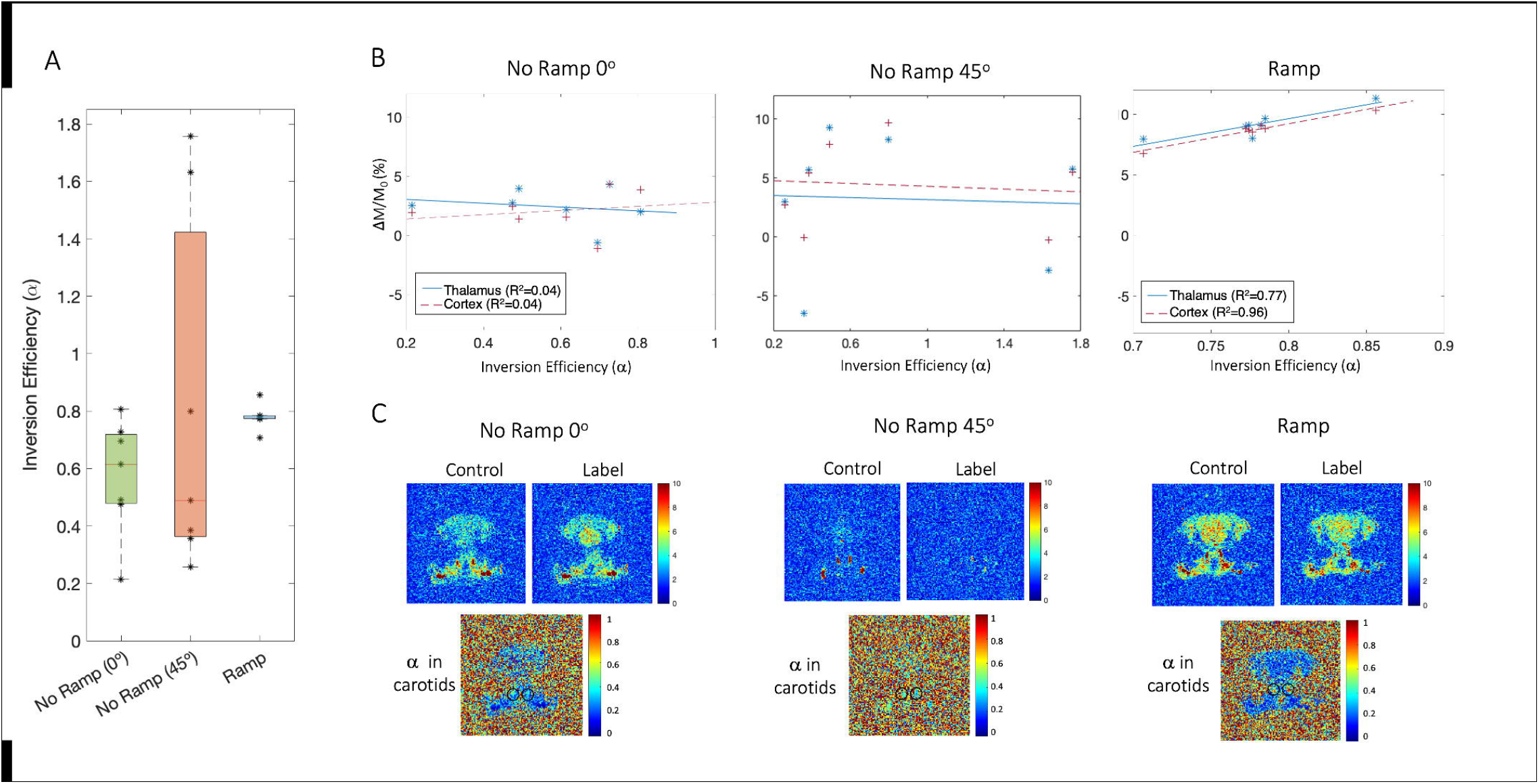
Inversion Efficiency calculation. (A) Inversion Efficiency (*α*) estimation for the 3 setups at the level of the carotids. Note that the average measured *α* for the *no ramp 45°* setup is higher than for the *ramp* setup since some of the measured values are higher than physiologically possible = 1. There is a dramatically smaller variance in the *ramp* compared to the other conditions. (B) Relationship between ΔM and across *α* setups in two specific ROIs (thalamic and cortical). Each dot represents one animal; (C) Magnitude images acquired with the fc-FLASH sequence. Note that in the *no ramp 45*^*o*^ setup, the carotids can hardly be spotted.

The relationship between rASL and *α* for the three scenarios is presented in Figure 2, panel B. The expected linear relationship - the more signal labelled (*α*), the more signal measured (*rASL*) - in the two ROIs chosen (R^2^ thalamus right side = 0.77 and R^2^ cortex right side= 0.96) was observed only for the *ramp* setup. Note the unphysiological negative ΔM/M_0_ values. Although it is not possible to have negative perfusion, in an imperfect experiment, the subtracted signals (ΔM) can yield negative rASL values, since the signal is probably not portraying actual perfusion. The labeling efficiency magnitude images acquired with the fc-FLASH sequence are shown in Figure 2, panel C, revealing that in the conventional *no ramp* 45 setup the carotids can hardly be spotted, while in the *ramp* setup there is a clear definition of the two circles representing the cross-section of the two carotids.

### 3.3. Mouse pCASL: CBF Maps

To illustrate the problem of between-animal variability, CBF maps of 4 representative mice are displayed in Figure 3 for the “conventional” *no ramp 0°* and the *no ramp 45°* setups, and at standard and high-resolution. Note that the low VS high resolution slices have different slice thickness and gap distance, so a more general comparison was performed since the anatomy is slightly different in the corresponding slices of the standard and high-resolution data sets. In mouse 1, the CBF maps show disparity between setups: the *no ramp 0°* presents extremely low CBF values brain-wide as well as artifacts on the more caudal slices, while the 45*°* tilt in the labeling plane generated physiologically relevant maps, although there is decay present in the more rostral slices.

In mouse 4, both setups show very low, unphysiological perfusion values. Mouse 6 displays asymmetry between the two hemispheres, in both setups. Finally, mouse 7 presents maps that resemble physiologically relevant CBF maps for both cases in the standard resolution images. However, in the high-resolution images, a slight asymmetry can be spotted in the more caudal slices.

**Figure 3.**
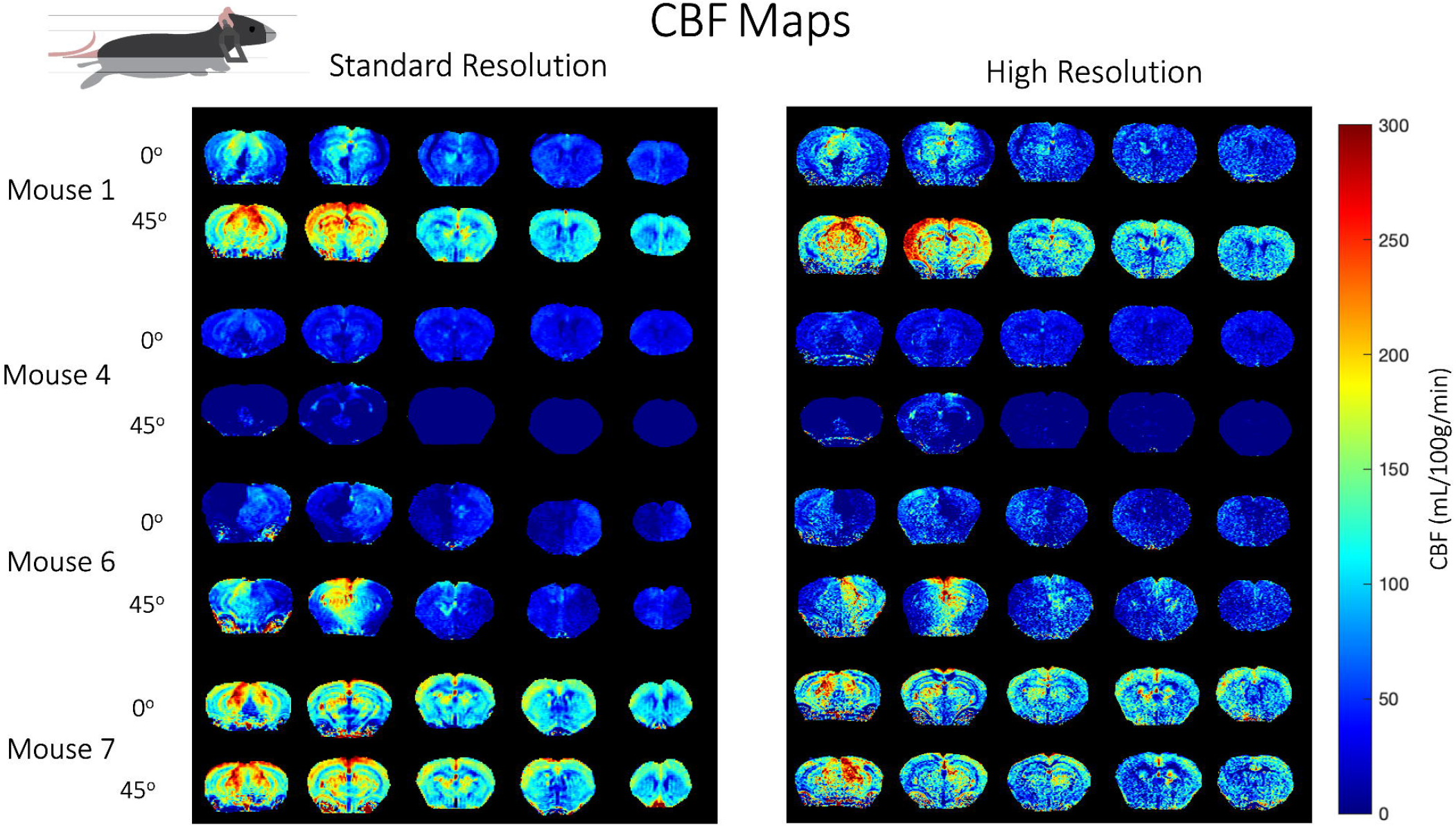
Quantitative CBF maps for standard and high-resolution acquisitions for 4 representative animals scanned with no *ramp* (*0°* and *45°*). There is high disparity between setups along with extremely low CBF values brain-wide and asymmetries across brain hemispheres.

CBF maps obtained for the N=6 mice scanned using the novel *ramp* setup, both at the standard resolution and at x15 higher resolution (when compared to state-of-the-art resolution reported in literature: 225x225x1500 μm^3^) ^9,10,16^, are presented in Figure 4. In contrast to the non-*ramp* setups, CBF maps are consistently of high quality, and present very little variability across animals. All maps reveal the expected CBF patterns in the healthy mouse brain, with greater perfusion in grey matter (e.g., cortex and thalamus) when compared to white matter. In the high-resolution images (100x100x500 *μ*m^3^), further structures become visible: perfusion in different cortical layers can be discerned, along with clearly delineated descending vessels. Even in the hippocampus, where CBF is lower compared with other grey matter regions, layers can be observed. Furthermore, the group analysis revealed that the *ramp* setup has very low IQR (Supplementary Figure 5).

**Figure 4.**
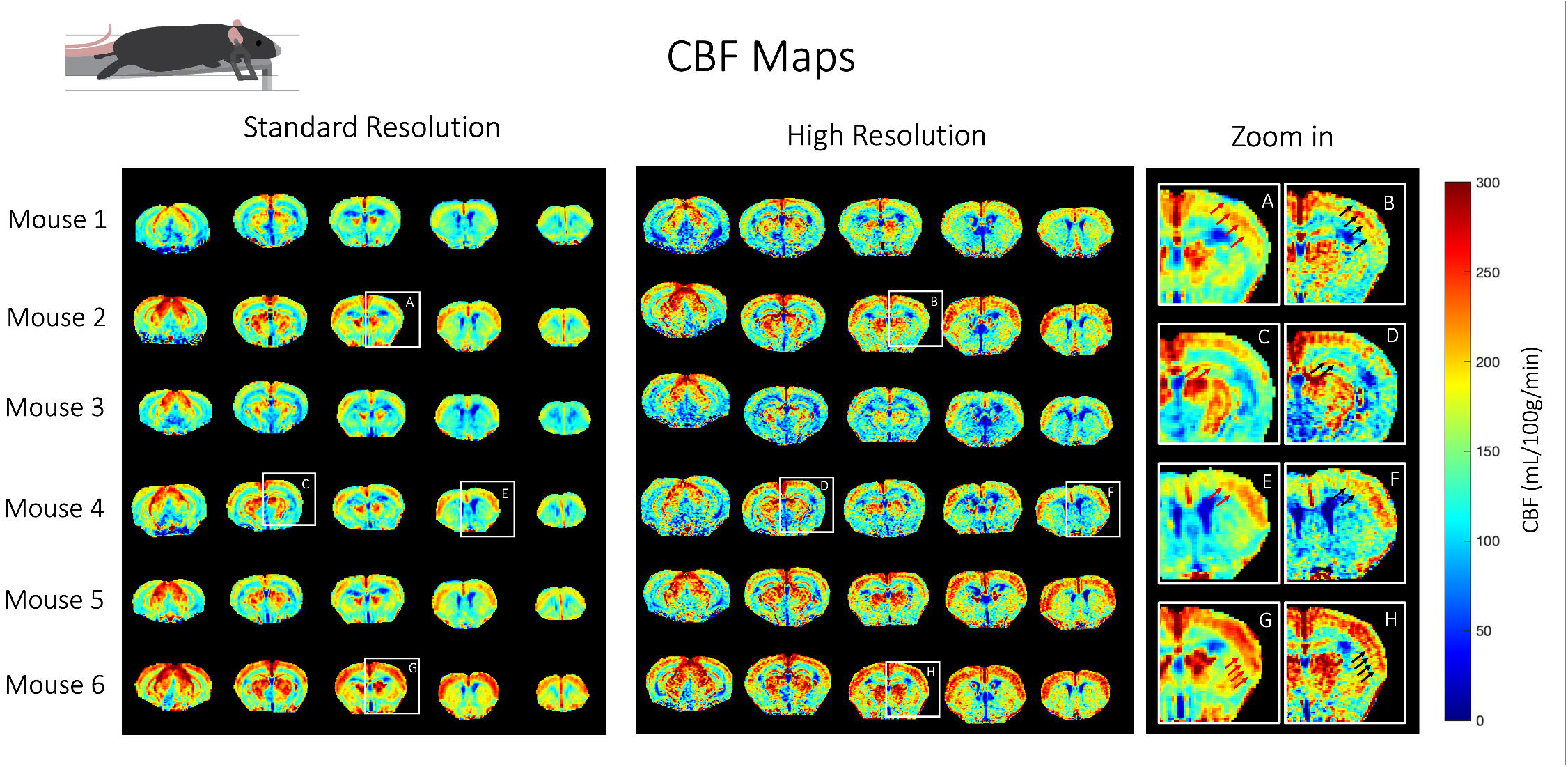
Quantitative CBF maps for standard and high-resolution acquisitions with the *ramp*. All maps show robust measures of CBF with very little variability across animals. The zoom-ins delimit findings that could only be attained through high-resolution imaging: (A,B) cortical layers; (C,D) hippocampal layers; (E,F) cortical local descending vessels; (G,H) columnar blood vessels. For each zoom-in, the structures that can be seen with the high resolution but not with the standard resolution are highlighted by back and red arrows, respectively.

### 3.4. Mouse pCASL: ROI Analysis

To further compare the 3 setups, we also performed an ROI analysis of rASL and CBF in two different regions: cortex and thalamus (right and left hemispheres) in the high-resolution scans; the results are shown in Figure 5. For both regions, the *ramp* setup gave the highest average value, the lowest standard deviation, and the lowest brain asymmetry across animals. The *no ramp 45°* setup presents highly dispersed values of both CBF and rASL across different animals and different regions of the brain, which highlights the inconsistency of the measurement. The *no ramp 0°* displayed lower rASL signal when compared to the *ramp* setup. When looking at inter-hemisphere differences (between symmetrical ROIs within the same animal), it is clear that the *ramp* setup presents, once again, the lowest asymmetries in both the CBF and rASL calculation when compared to other setups.

**Figure 5.**
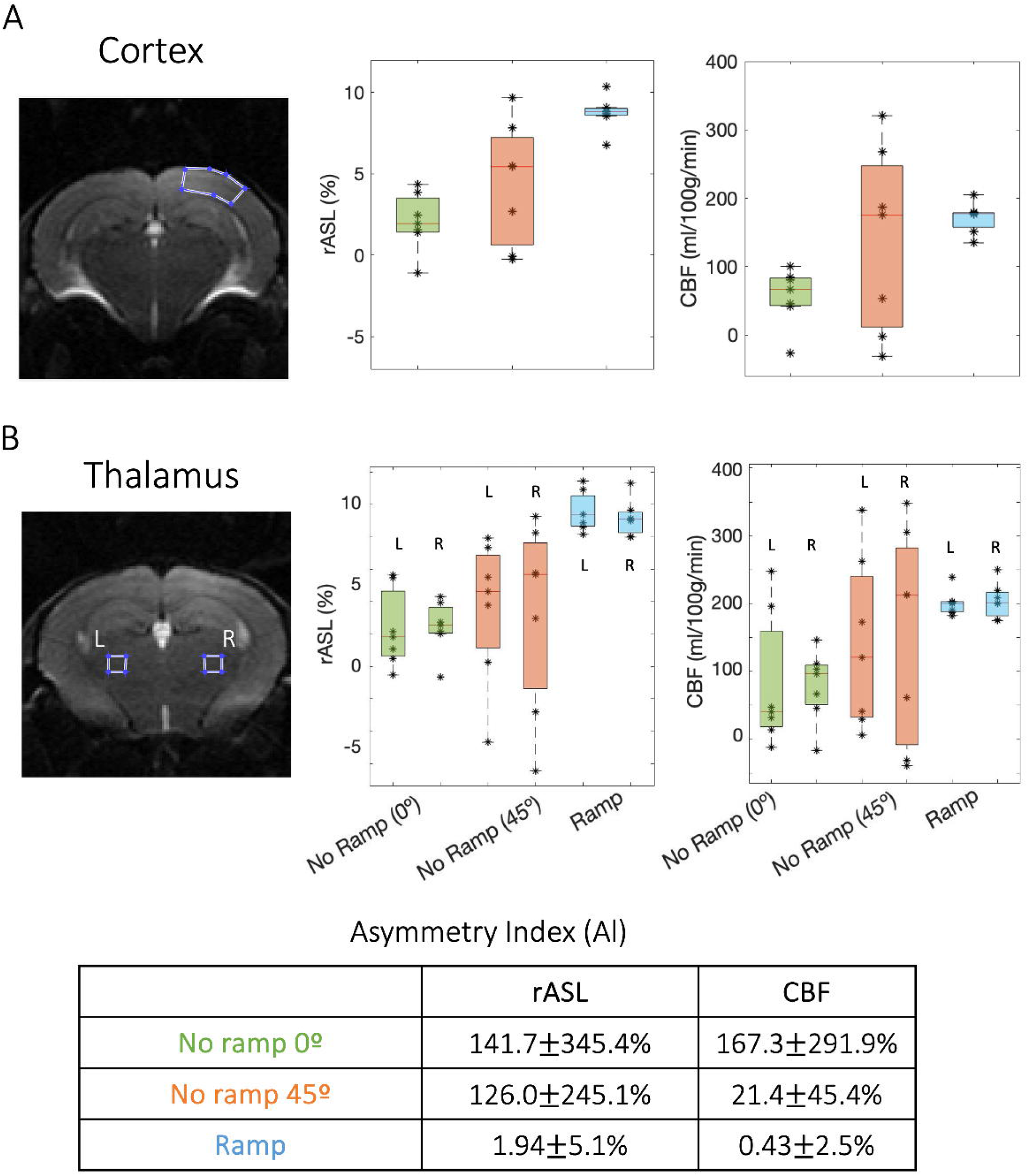
Comparison of *rASL* and *CBF* in three ROIs between setups. (A) cortical ROI and B) thalamic ROIs; Across all the ROIs, the *ramp* setup shows the lower standard deviation across animals and the highest *rASL*. In thalamus, the *ramp* setup presents the lower brain asymmetries. Note that the *no ramp 45°* setup presents highly dispersed values of both *CBF* and *rASL* across different animals and different regions of the brain, which depicts the inconsistencies of the measurement.

### 3.5. Mouse pCASL: stroke model

To further assess the validity of our methodology, we also applied it to a mouse model of photothrombotic stroke. Longitudinal images for a representative stroked mouse at 0h, 1d and 5d after stroke are shown in Figure 6. At the earliest time point (t= 0h), the left cortex shows a severe reduction in CBF, with strong hypoperfusion near the infarct core. The core is much better delineated in the high-resolution images, as well as the descending cortical vessels that are clearly outlined in the hemisphere contralateral to the stroke. After 24 hours, part of the cortex has already been reperfused and the infarct core size increased. The high-resolution images reveal interesting new information: this enlarged core contains vessels where very little CBF can be measured (<50 ml/100g/min) (Figure 6B). This information could not be retrieved from the standard resolution images. Finally, after 5 days, the stroke core has reduced in size, showing almost no perfusion in the affected area. In this more chronic stage, a proper quantification of the lesion volume would be much better achieved with the use of the high-resolution images, where the borders of the stroke core are clearly defined.

**Figure 6.**
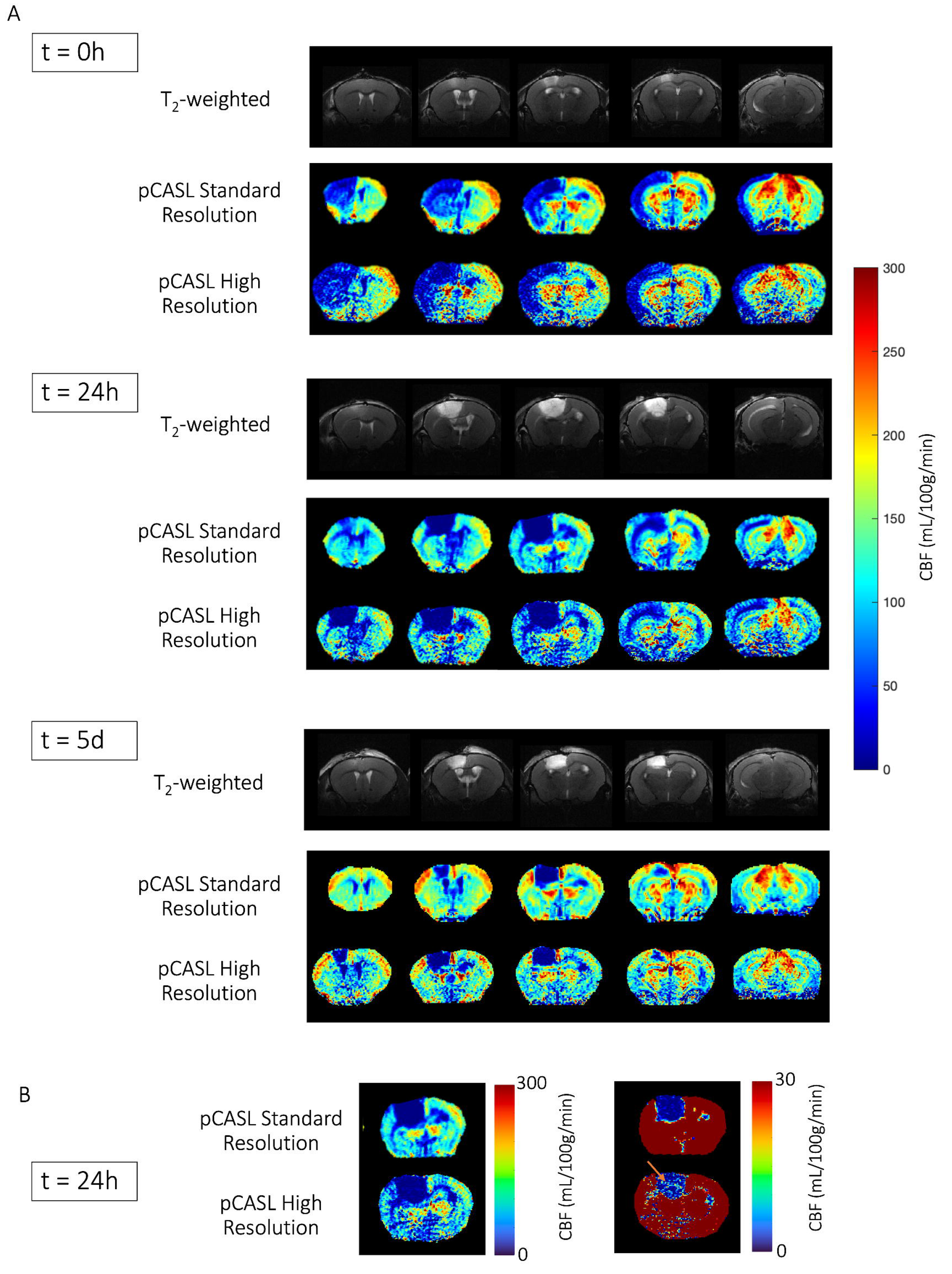
A) CBF maps of a representative mouse induced with photothrombotic stroke, scanned at 3 different timepoints with the *ramp* setup. Visible differences can be observed from a more acute stage (where the lesion is vast and comprises most of the cortex in the hemisphere where the lesion was induced) to a more chronic stage (where the core of the infarct is severely hypoperfused, mostly containing dead tissue). The high resolution data yielded improved discrimination at border zones, namely at the stroke core, due to the decrease in partial volume effects. Note that the low VS high resolution slices have different slice thickness. B) Highlight of the perfusion inside the stroke core at t=24h for slice 3. The remaining vessels seen in the high resolution scans are highlighted by the orange arrow.

### 3.6. Rat pCASL

We also sought to increase the resolution in rat pCASL imaging. This implementation did not require any additional setup due to the larger anatomy of rats (when compared to mice) that avoids the use of ear bars (which alter the optimal positioning of the carotids). Figure 7 shows CBF maps at standard resolution (234x234x1000 *μ*m^3^, top) and the x6 higher resolution (110x110x750 *μ*m^3^, bottom) in 3 control rats and 1 representative rat where a stroke was induced. High quality CBF maps were obtained in all the animals with the expected CBF patterns in the brain, namely greater perfusion in grey matter (e.g., cortex and thalamus) when compared to white matter. These maps show high reproducibility across different rats, for both standard and high-resolution settings. The average α of 87.0±4.0% was also highly consistent. Note that in the x6 higher resolution images, small structures can be discerned more clearly than in the standard resolution images.

**Figure 7.**
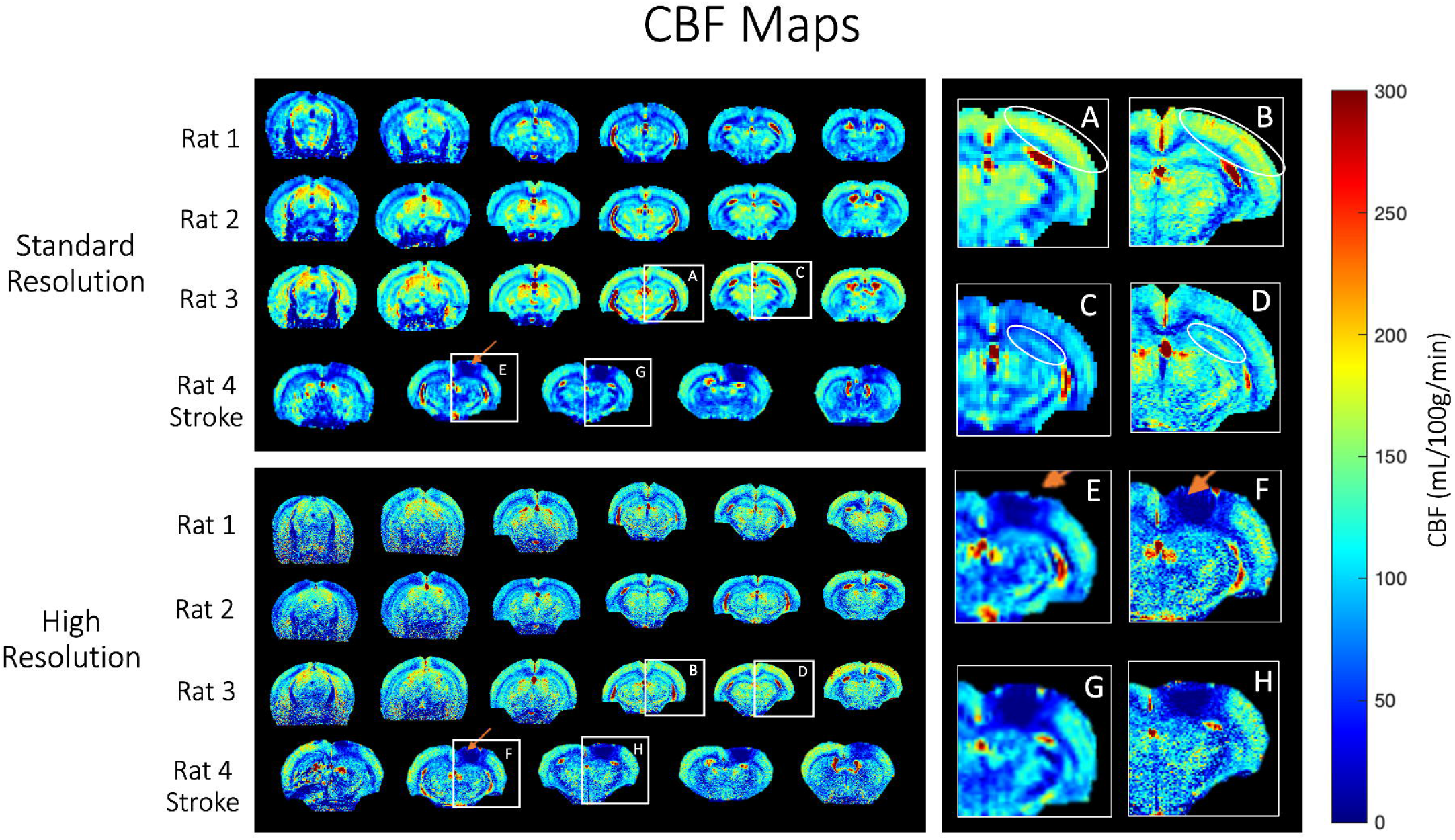
Quantitative CBF maps for standard and high-resolution acquisitions across 3 healthy rats (6 slices per animal). In rat 4, a stroke was induced 24h prior to the acquisition. The orange arrow indicates the stroke core. The zoom-ins delimit findings that could only be attained through high-resolution imaging: (A,B) cortical layers and local descending vessels; (C,D) hippocampal layers; (E,F & G,H) proper delimitation of the stroke core.

As a proof-of-concept in rats, a photothrombotic stroke was induced in one rat and the animal was scanned 24h post-stroke induction (Figure 7). The core is visibly hypoperfused and regions of less perfusion around the core – penumbra – can be seen but only in the high-resolution images. Additional details regarding the volume of the stroke core can be spotted in the high-resolution images as outlined by the zoom-in presented in Figure 7.

## 4. Discussion

Rodent models play a critical role for characterising biological processes in health and disease and are important tools in neuroscience and biomedicine research.^33^ Given the importance of CBF at specific laminar and columnar levels, high-resolution in-vivo pCASL imaging, with its capacity for longitudinal studies and non-invasive access to the brain, can play a decisive role in pre-clinical neuroimaging. In this work, we set out to improve the state-of-the-art resolution by over an order of magnitude in mice (and by a factor of at least 6 in rats) and to obtain high reproducibility of CBF maps across animals. We found that, in mice, specific anatomical constraints involving the relative position of carotids and brain under conventional settings in the MRI scanner generate highly variable and limited pCASL labeling efficiency. We overcame this limitation by introducing a novel setup that stabilizes mouse positioning for ASL scanning and showed that it produced consistently high inversion efficiency in mice, dramatically reducing between-animals variability. The use of a cryogenic coil^24^ significantly enhanced the sensitivity of the pCASL measurements to the point where the order-of-magnitude increase in spatial resolution was achievable. Our CBF measurements aligned with the values reported in the literature for both rats and mice, showing that rats have lower CBF compared to mice.^10,34,35^ An initial application to a mouse (and rat) model of stroke demonstrated the validity of the proposed method to detect the expected alterations in CBF in such pathological conditions. Below, we discuss each of these aspects.

### 4.1 New experimental setup for mice pCASL imaging

High spatial resolution is crucial to obtain a detailed representation of small brain structures in perfusion images. To achieve this goal, we made use of the pCASL sequence developed by Hirschler et al. (2018)^15^, combined with a cryogenic coil to enhance the sensitivity of the measurement in small animals.^24^ The implementation of this sequence alone accounts for B_0_ and B_1_ inhomogeneities at the labeling place, and the cryoprobe has the potential to enhance the SNR, but as shown, these two improvements are insufficient to achieve high-resolution scans in mice (although it can be more easily achieved in rats). Indeed, the typical positioning of mice in the scanner - head fixed with ear bars and mouth fixed by the upper incisors locked on to a bite bar – bends the carotids relative to the brain, particularly when using a cryogenic coil. In this case, the mouse head is always elevated with respect to the body to maximize the proximity to the coil, thereby leveraging the maximum SNR gain offered by the cryogenic coil. This in turn results in an abnormal positioning of the carotids relative to the labeling plane, leading to suboptimal flow-induced adiabatic inversion and therefore, impaired labeling efficiency in pCASL.

Our custom-built *ramp* was able to stabilize the mouse positioning while keeping the head close to the coil for SNR purposes, as shown in Figure 1B. This obviates nearly all of the complications arising from the carotid angle and enables the reliable CBF quantification with high resolution. Indeed, it is notable that the variability caused by the inconsistencies in the labeling efficiency from both the no *ramp* setups (Figure 3) – i.e., non-physiological effects – is much larger than the underlying physiological variability in blood flow across animals (Figure 4). We also observed that there is systematically greater SNR in the *ramp* setup when in comparison to other setups (Supplementary Figure 3). Even though in both scenarios the mouse head is positioned as close as possible to the coil, we believe that the *ramp* causes a subtle change in position in the more caudal slices (in which the ROI analysis was performed), making the animal slightly higher and therefore closer to the coil – another advantage of using the *ramp*. We note that in rats, the positioning within a typical cryoprobe bed does not require the use of ear bars and, hence, their heads are not elevated in relation to the body, obviating the use of such a *ramp* for rats.

We also tested whether the application of our *ramp* design could assist in enhancing the quality in the more conventional non-cryogenic coil setup (Supplementary Figure 6). Our results suggest that with standard resolution the *ramp* setup indeed improves the reproducibility of the CBF maps. However, as expected from the lower SNR of room-temperature probes, the higher-resolution data was of poor quality, suggesting the importance of the cryogenic coil for high fidelity CBF mapping. We further note that the *ramp* setup was optimized specifically for the cryogenic coil and due to differences in mouse bed design between the room temperature setup and cryogenic setup, there is likely room for further optimization and improvement for the *ramp*.

### 4.2 Reliable CBF quantification

To investigate the effectiveness of the proposed setup for pCASL perfusion imaging in mice, we tested three different combinations of labeling plane positioning with and without the use of the *ramp*. In the no *ramp 0°* setup, there is a 90° angle between the carotids and the brain (in the sagittal plane), so a labeling plane that is parallel to the imaging slices in the brain is not expected to label the carotids properly (Figure 1 C, Left). On the other hand, in the *no ramp 45°* setup, the carotid and vertebral arteries would be aligned to the labeling plane due to the tilt angle (Figure 1 C, Left). However, this combination also resulted in high variability in labeling efficiency across animals, likely due to subtle idiosyncratic positioning of the arteries in the animals and out-of-plane tilts, leading to a suboptimal flow-induced adiabatic inversion. Another possible cause for the inconsistencies arising from the use of this setup is the likelihood that the tilt in the labeling plane will have an effect in the imaging region, due to the proximity between the two caused by the sharp angle prescribed. It is also interesting to highlight that the α values for the no *ramp* 45 degree setup were observed to occasionally go beyond what is physiologically possible (α>1), i.e., 100% inversion of spins flowing through the carotids (α=1). This once more underlines the inconsistency of the measurement when the conditions are suboptimal.

Indeed, after quantification, the CBF maps obtained with both no *ramp* approaches do not reveal the expected patterns in perfusion maps, have low reproducibility across animals and present asymmetries within brain hemispheres in the same animal. These observations are in agreement with the ROI analyses, which revealed lower CBF values for the *no ramp 0°* setup in the two brain regions analysed, reflecting the less successful labeling caused by the variable/incorrect positioning of the animal and hence the labeling plane. The *45° ramp* setup shows high variability across mice, which reflects the measurement inconsistencies. These discrepancies may be caused by the subtle differences in positioning across different animals (or even the same animal when scanned at different timepoints). We note that rarely, the correct labeling plane positioning can be achieved using the tilt, but we stress that we were not able to predict the conditions in which this occurs. Furthermore, the hemisphere asymmetry in both *no ramp* setups could be related with a less successful interpulse phase correction, promoted by the inconsistent positioning of the labeling plane in relation to the feeding arteries of the brain.^15^

All these inconsistencies vanish with the *ramp* setup introduced here. Hemispheric brain asymmetries were not detected in any of the animals scanned with the *ramp*, and we could consistently obtain perfusion maps for all animals within the plausible and expected physiological range (∼200 ml/100g/min).^36^ Furthermore, the *ramp* setup was the only configuration in which high labeling efficiency was consistently achieved in all animals, and the expected linear relationship between this and the ASL signal was observed. The other two configurations show only a random distribution of values across the two different brain regions evaluated, which reveals an inefficient labeling and/or an unsuccessful quantification of the labeling efficiency. Although the average α measured for the *no ramp 45*^*0*^ setup (0.80±0.63) was higher than for the *ramp* (0.78±0.04), it exhibited a very large variability. In fact, some of the values measured with the *no ramp 45°* setup are implausible (higher than =1, which corresponds to labeling 100% of the spins flowing through the labeling plane). This unveils, once more, the inconsistency of the measurement and the difficulties in properly quantifying the labeling efficiency itself when the setup is not optimized.

In the photothrombotic model animals scanned with the *ramp*, our findings show small variability among animals and are consistent with CBF flow alterations in stroke models reported in literature (Supplementary Figure 7).^37^ Indeed, visible differences can be observed from a more acute stage (where the lesion is vast and comprises most of the cortex in the hemisphere where the lesion was induced) to a more chronic stage (where the core of the infarct is severely hypoperfused, mostly containing dead tissue).^37^

### 4.3 High-resolution pCASL imaging

The previous studies harnessing pCASL in mice made use of a transmit-receive volume coil and implemented a higher number of label-control repetitions (60 repetitions instead of the 30 used in this study, which requires double the acquisition time).^9,10,16^ The achieved spatial resolution (225 × 225 *μ*m^2^; slice thickness: 1.5 mm) was much lower than the lowest resolution reported in this study for mice (146x146 *μ*m^2^, slice thickness=1 mm). The high resolution presented here for the first time (100x100 *μ*m^2^, slice thickness= 0.5 mm) is a significant improvement in the state-of-the-art resolution of pCASL mouse images, raising the bar by a factor of 15. The key factor in this improvement in spatial resolution is the cryogenic coil, which significantly reduces noise and leads to a higher SNR.^24^

High resolution CBF images may provide relevant insights on blood supply to the brain and be able to distinguish changes at both columnar and laminar levels.^37–39^ Indeed, in the high-resolution images reported here both for mice (Figure 4) and rats (Figure 7) the columnar cortical pattern of most animals could be observed, which is consistent with the descending arterioles and ascending venules anatomy, as well as with cortical layering reflecting different vascular densities.^40,41^ In the hippocampus, different layers also become visible with the high-resolution scans as well as specific vessels that perfuse more ventral areas. Furthermore, in the animal models of stroke here presented, the high-resolution data yielded improved discrimination at border zones, namely at the stroke core, due to the decrease in partial volume effects that is observed when acquiring smaller voxels. For instance, in the rat high-resolution stroke images (Figure 7 – zoom in) there is a clear definition of the stroke core as well as the surrounding penumbra – tissue at risk but not yet irreversibly damaged – that is crucial information when evaluating/assessing the progression of the disease and plays an important role on the anticipation of the outcome.

### 4.4 Limitations

As in all studies, several limitations can be identified in our work. One possible limitation of this study is that the mice used were relatively young and light. However, we have already applied this setup (*ramp* with the exact same height) to a Parkinson’s disease study and scanned animals at 39±3 weeks of age and weighing 42g±15 g.^42^ The results were sustained in these conditions. Hence, there is no problem in using our setup for mice at least up to ∼40 gr. Regarding the rats, no additional setups were used so scanning larger animals would not be a problem as long as they fit inside the scanner – the maximum weight that we typically use inside the scanner bore (20 cm) is ∼ 550 g.

Another important technical limitation in mice is that we performed single-slice acquisitions. Although multi-slice acquisitions would be possible with this sequence, extending the acquisitions to more than one slice entails undesirable effects that would influence our CBF estimation, including saturation effects due to the proximity of the slices ^43^ This could be solved in the future by using a 3D readout or by developing specific corrections/theories for multi-slice pCASL acquisitions or by using simultaneous multi-slice (SMS) acquisition schemes that have been already presented for rodents.^44^ Furthermore, it should be mentioned that the typical pCASL sequence can be especially vulnerable to the outflow effect in areas with high flow rates due to its lengthy labeling duration (∼3 seconds), which can cause CBF to be underestimated. This limitation could be overcome by implementing a multi delay strategy that accounts for differences in arterial transit times (ATT).^9,45–47^ This would also allow for the estimation of other perfusion related parameters (including the ATT) that may be altered in pathologies such as the ischemic stroke. In principle, higher resolution imaging may be possible this way using novel denoising approaches that make use of the information sparsity in such measurements.^48–52^

Another limitation for some applications involves our use of isoflurane anaesthesia. In functional MRI for example, CBF measurements could be important but hampered by isoflurane’s uncoupling of neurovascular coupling.^39,53^ More generally, isoflurane has a known vasodilatory effect^54^ that will eventually alter the brain haemodynamics in rodents in comparison to their natural awake state.^10^ In the future, awake mice could be used to avoid any confounding effects from anaesthesia ^10,55,56^

One drawback of our single-shot EPI approach is the longer TE compared to previous studies^15^; still, we observed highly consistent results across animals and high relative ASL signal, suggesting that this was not a major confounding factor. In the future, the resolution could be further increased by acquiring EPIs with multiple segments, or faster spiral trajectories.

Finally, we note that no denoising strategies were implemented here, but approaches such as non-local-means^51^ or MPPCA^48,49,52^ could further augment the SNR gains reported here, leading to even higher resolution and therefore serve well for high-definition perfusion imaging in the healthy and diseased rodent brain.

## 5. Conclusions

Using a cryogenic coil and a novel *ramp* setup for proper mouse carotid positioning, we achieved an order-of-magnitude improvement in pCASL spatial resolution without increase in scan time, as well as much higher reproducibility of CBF measurements compared to state-of-the-art methods. We applied our novel setup and showed its potential for detecting penumbra in stroked rats and mice. Our results bode well for future applications of quantitative CBF mapping in preclinical neuroimaging.

## List of abbreviations

α: inversion efficiency
AI: asymmetry index
ASL: arterial spin labelling
ATT: arterial transit time
BBB: blood brain barrier
CASL: continuous arterial labelling
CBF: cerebral blood flow
EPI: echo planar imaging
fc-FLASH: flow-compensated fast low angle shot
LD: labelling duration
MRI: magnetic resonance imaging
PASL: pulsed arterial spin labelling
pCASL: pseudo continuous arterial labelling
PLD: post-labelling delay
RF: radiofrequency
rASL: relative ASL signal
ROI: region of interest
S1: somatosensory cortex
SAR: specific absorption rate
SNR: signal to noise ratio
TE: echo time
TOF: time-of-flight
TR: repetition time
IQR: inter quartile range

## Acknowledgments

The authors acknowledge the vivarium of the Champalimaud Centre for the Unknown, a facility of CONGENTO which is a research infrastructure co-financed by Lisboa Regional Operational Programme (Lisboa 2020), under the PORTUGAL 2020 Partnership Agreement through the European Regional Development Fund (ERDF) and the Champalimaud Communication, Events and Outreach team. We would also like to acknowledge Fundação para a Ciência e Tecnologia (Portugal), under project LISBOA-01-0145-FEDER-022170, grant 2021.08457.BD and LARSyS funding (DOI: 10.54499/LA/P/0083/2020, 10.54499/UIDP/50009/2020, and 10.54499/UIDB/50009/2020).

## Author contribution statement

N.S., P.F. and S.P.M. designed the project and the necessary experiments, and N.S. and P.F. oversaw the implementation of the project. S.P.M. acquired and analysed the data. N.S., P.F. and S.P.M. wrote the paper. L.H. and E.L.B. provided the pCASL sequence. All authors discussed the results and implications and commented on the manuscript at all stages.

## Notes

### Competing Interest Statement

The authors have declared no competing interest.

### Summary of Updates

Most sections were altered as well as the figures since the paper underwent an extensive revision before publication in a peer-reviewed journal.

