## Supplementary information for "High-resolution perfusion imaging in rodents using pCASL at 9.4 T"

**Supplementary Figures**

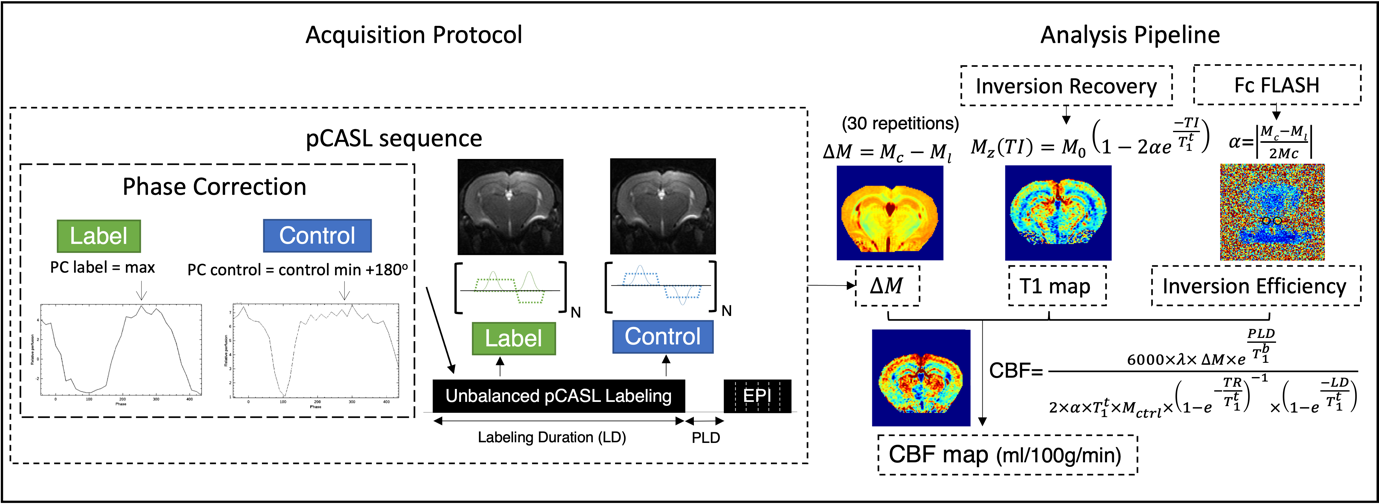
**Supplementary Figure 1.** (Left) Schematics of the acquisition protocol displaying the label and control phase corrections implemented and the pCASL sequence. (Right) Analysis pipeline with different inputs for CBF quantification and representative images for each step. $\Delta$M is the signal difference between control and label acquisitions averaged over 30 repetitions.

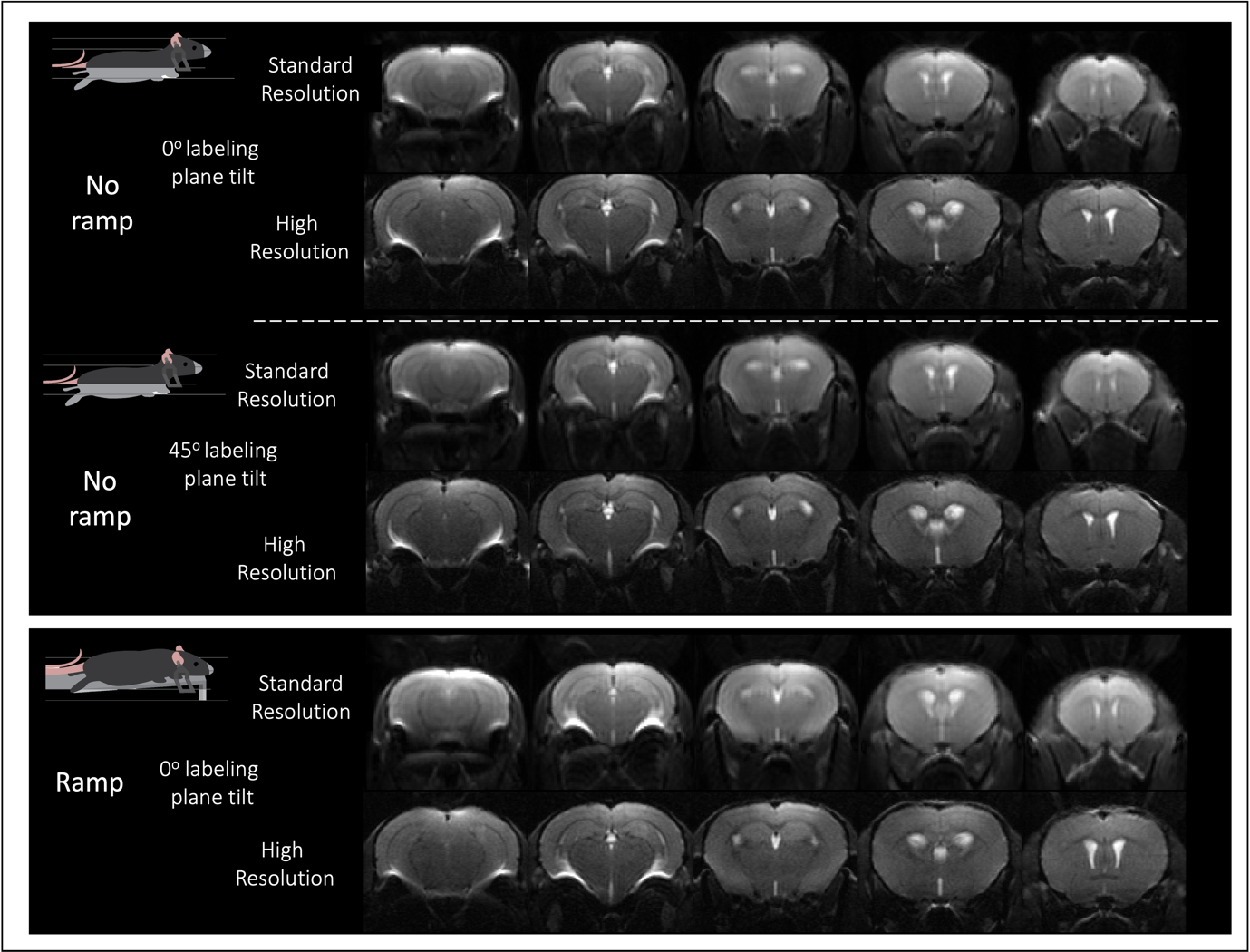

**Supplementary Figure 2.** Representative raw data acquired with the 3 setups (*no ramp 0^o^, no ramp 45^o^, ramp*) at standard and high resolution in mouse. In all cases, the raw data are consistent and with high quality, revealing that all the EPI acquisitions were properly achieved and did not vary within setups.

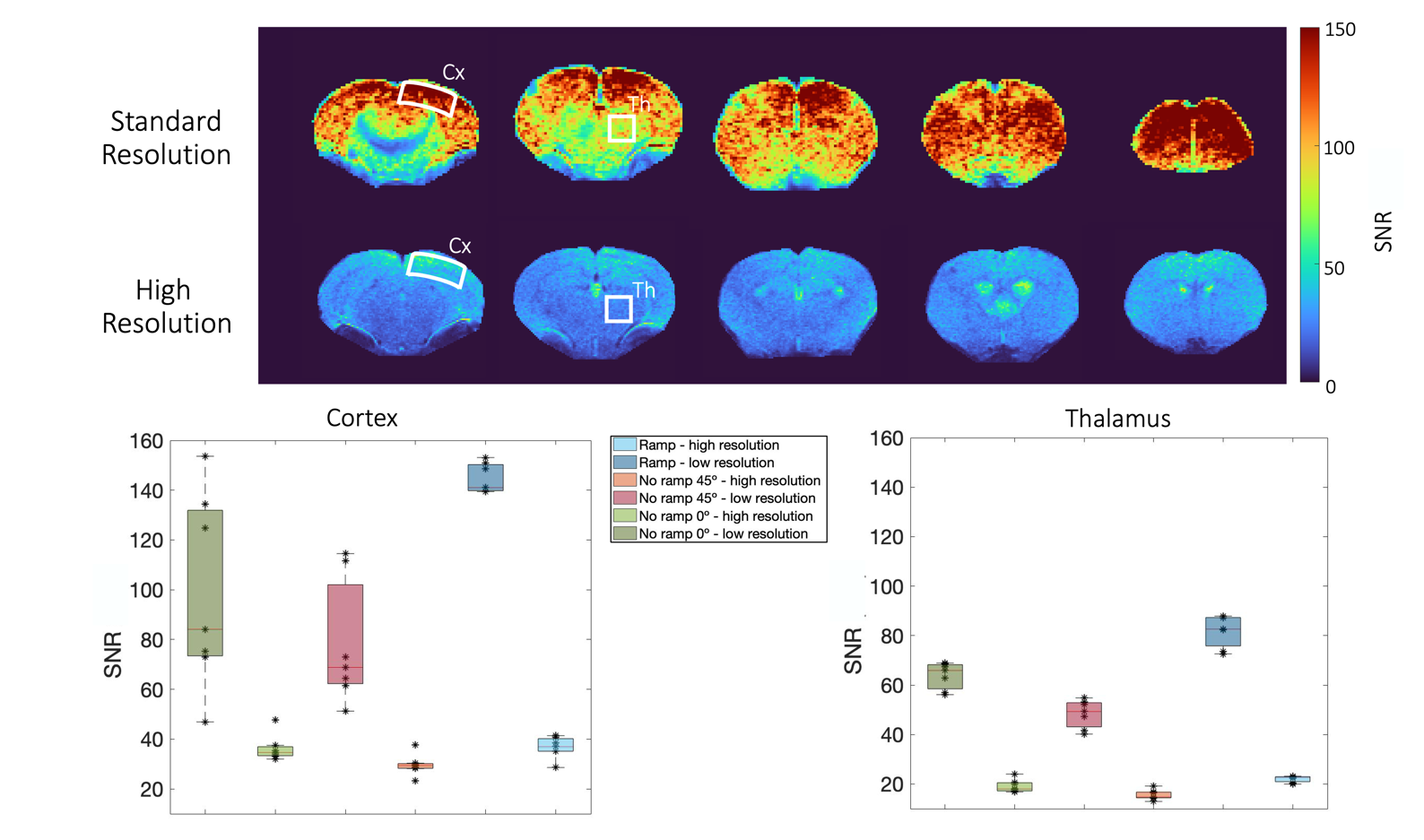
**Supplementary Figure 3.** (Top) SNR maps of the control images for one representative animal scanned with the ramp: average of 30 repetitions for 6 slices. (Bottom) Comparison of SNR between standard and high-resolution acquisitions in control images acquired with the 3 setups (no ramp 0^o^, no ramp 45^o^, ramp) in two ROIs (cortex and thalamus).

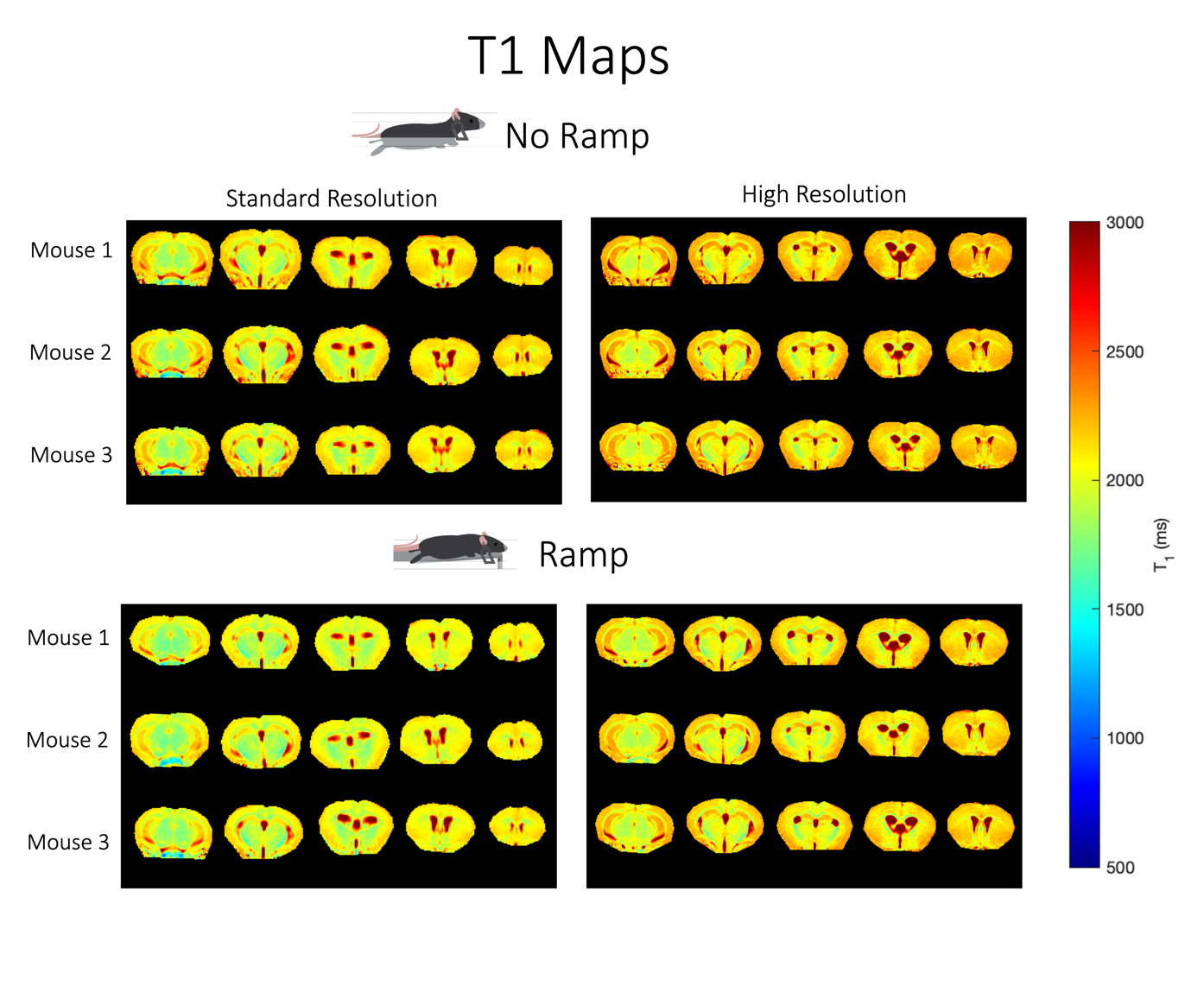
**Supplementary Figure 4.** Quantitative T1 maps for standard and high-resolution acquisitions for 3 representative animals scanned with the ramp and for 3 representative animals scanned with *no ramp* (*0^o^* and *45^o^*).

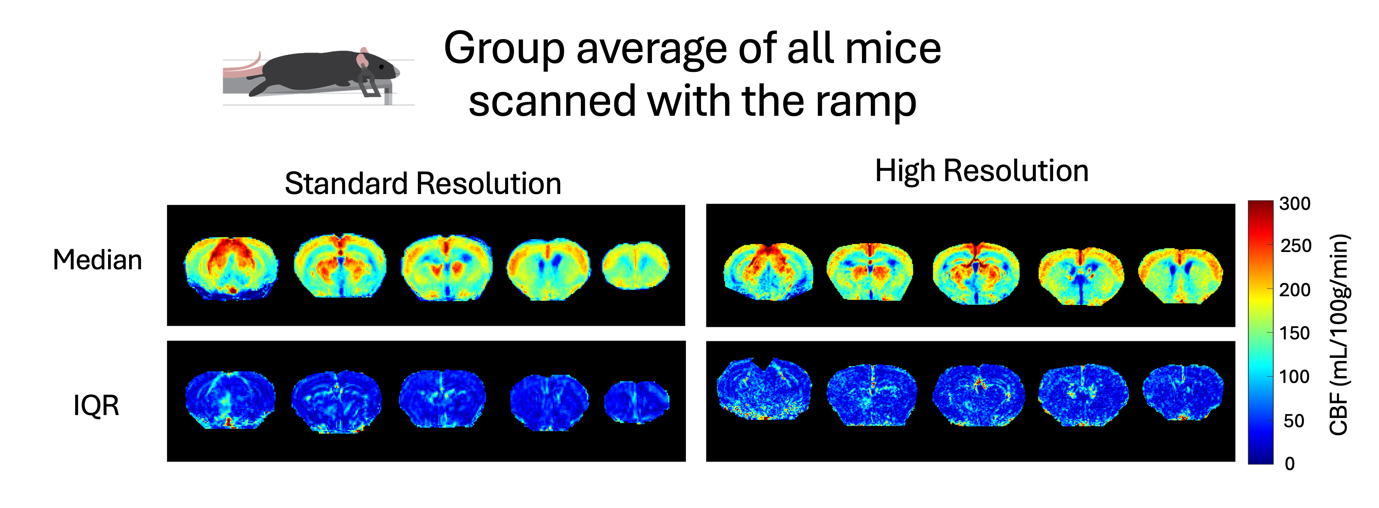

**Supplementary Figure 5.** Group average of all mice scanned with the ramp (N=6). Top: Median; Bottom: Interquartile range (IQR). Note the small variability for the CBF maps across animals.

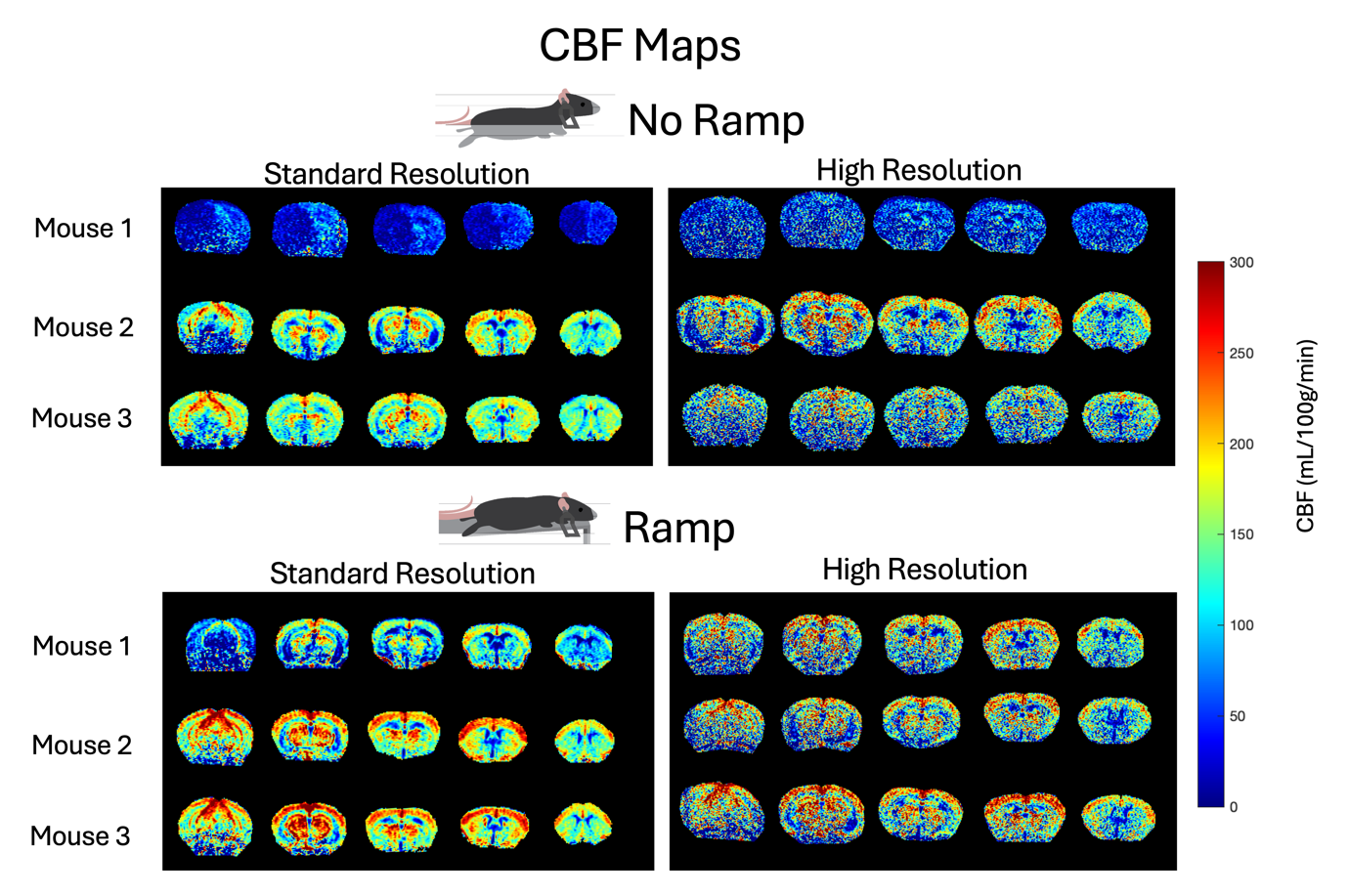

**Supplementary Figure 6.** Quantitative CBF maps for standard and high-resolution acquisitions for 3 animals scanned with no ramp 0^o^ and with the ramp using a room-temperature coil. A high variability is observed when the ramp is not used, (note that CBF in the first mouse is very different from the other two), which is remedied when the ramp is used, suggesting the usefulness of the ramp even under room temperature coil settings. However, we note that higher resolution images fail to produce high quality data (as expected due to the room temperature coil’s lower SNR); still, the standard resolution provides acceptable results.

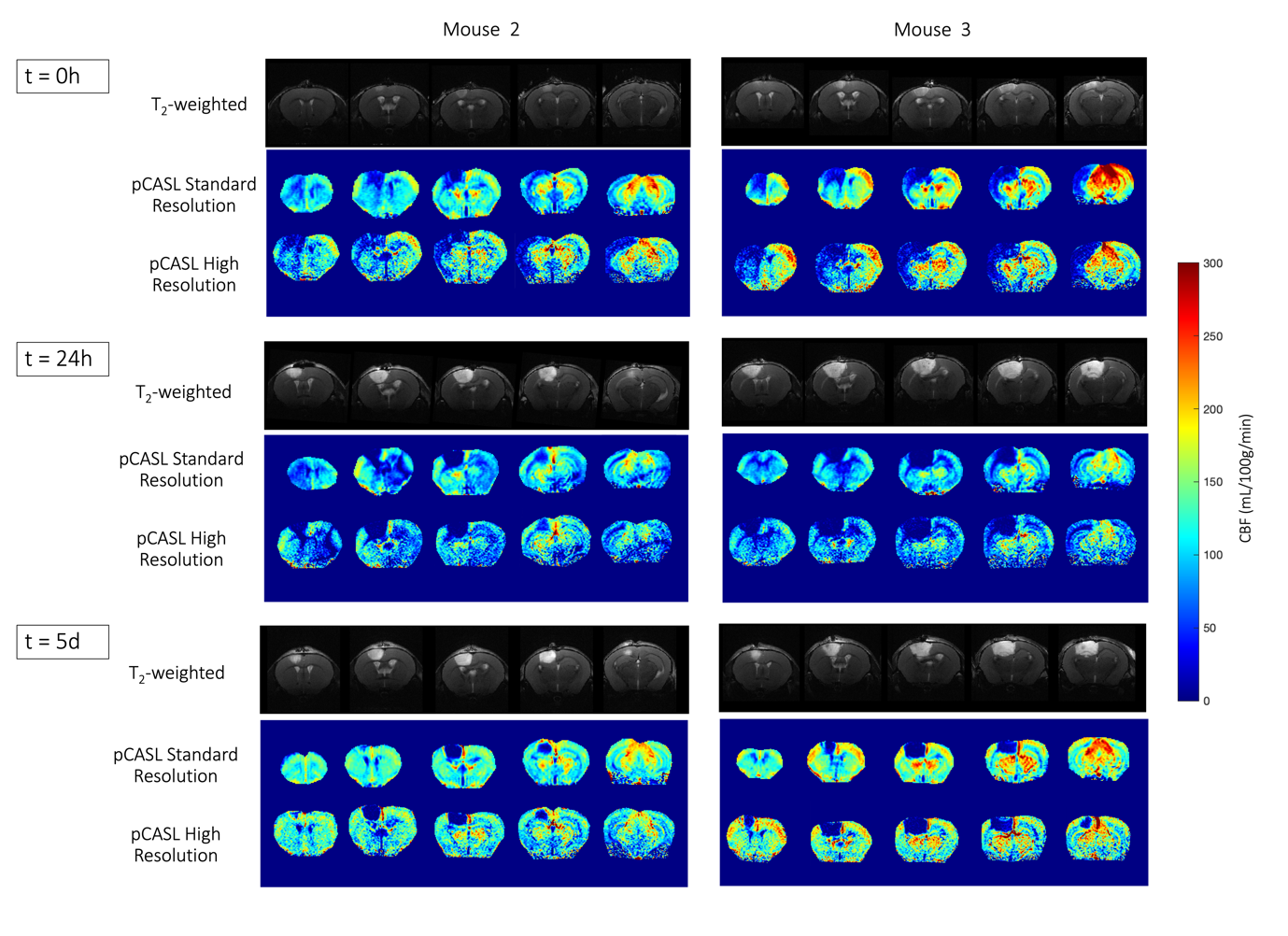

**Supplementary Figure 7.** CBF maps of two mice induced with photothrombotic stroke, scanned at 3 different timepoints with the ramp setup. Visible differences can be observed from a more acute stage (where the lesion is vast and comprises most of the cortex in the hemisphere where the lesion was induced) to a more chronic stage (where the core of the infarct is severely hypoperfused, mostly containing dead tissue). The high-resolution data yielded improved discrimination at border zones, namely at the stroke core, due to the decrease in partial volume effects. Note that the low VS high resolution slices have different slice thickness.
